# CRISPR-CAS9 MEDIATED EDITING OF THE *LACZ* GENE FOR OPTIMIZING LIPASE PRODUCTION AND CHARACTERIZATION IN ENGINEERED *ESCHERICHIA COLI* USING POTATO PEEL AS A GLUCOSE SOURCE

**DOI:** 10.1101/2024.09.02.608350

**Authors:** Joseph Bamidele Minari, Idowu Samuel Dada, Dhikrullah Oluwatope Abdulazeez, Gift E. Nwosu

## Abstract

Improper disposal of potato peels harms the environment, wasting nutrient-rich resources that could be used for beneficial enzyme production like lipase. This study aimed to edit the *lacZ gene* in *Escherichia* coli using utilizing the CRISPR Cas9 technology. Both edited and unedited *E. coli* were used for submerged fermentation of potato peels to produce and characterize lipase. The *lacZ* gene was edited using the CRISPR Cas9 technology, and the efficiency was measured using multiplex PCR and gel electrophoresis. To measure lipolytic activity, olive oil screening was performed. The temperature and pH of the lipase were used to characterize it after partial purification and submerged fermentation at 10°C, 30°C, and 45°C. Blue colonies indicated the *lacZ gene* was unedited, *lacZ* gene editing and repair was demonstrated by white colonies, and no colonies demonstrated the editing but not repair of the *lacZ* gene. Bands at 1,100 bp indicated unedited *lacZ gene*, while 650 bp showed edited *lacZ gene*. Increased cell mass was observed at 10°C. CRISPR Cas9 edited *E. coli* showed a clearer zone than the unedited in the lipase medium. The highest lipase activity from both edited and unedited *E. coli* was at 35°C, and the lowest at 65°C. Optimal pH was 7, with lowest activity at pH 4. The CRISPR Cas9 edited *E. coli* demonstrated significantly higher enzyme activities (p<0.05). This study concluded that CRISPR Cas9-mediated *lacZ gene* editing in *E. coli* enhances its ability to utilize potato peels, increasing lipase production.

## 1.0 Introduction

The food processing industry produces potato peels as a byproduct, which can be used as a valuable, cheap, and affordable starting material for value addition, product extraction, and the synthesis of commercially significant compounds including dietary fiber, biopolymers, natural antioxidants, and enzymes [1]. Production of enzyme has attracted both academics and industry since its inception. In the past, scientists have identified various enzymes and investigated their catalytic properties to comprehend intricate biological processes [2], consequently, to directly manage or make use of them [3]. One of the greatest examples of enzymes with particular substrate selectivity is the lipase (triacylglycerol hydrolase, EC 3.1.1.3), which is a member of the esterase family (EC 3.1.1.X) [4]. Lipases are enzymes that catalyze the aggregate or fractional hydrolysis of fats and oils, resulting in the release of free unsaturated fats, diacylglycerols, monoacylglycerols, and glycerols [5]. In comparison to lipases derived from plants and animals, microbial lipases are more useful due to their high yield, ease of genetic modification, diversity of catalytic activities, adaptation to different environmental conditions, consistency over time because there are no seasonal fluctuations, and rapid rates of microorganism growth in comparably less expensive medium [6]. Bacterial and fungal sources are the most often used microorganisms in the industry for the synthesis of lipases [7]. The most significant genera that have been utilized among the extracellular lipase-producing bacteria are *Archromobacter*, *Arthrobacter*, *Staphylococcus, Alcaligenes, Bacillus, Micrococcus, Lactobacillus, Burkholderia, Chromobacterium, Bacillus, Propionibacterium, Escherichia* [7]. *Escherichia coli* is one of the most often utilized cellular factories for the synthesis of biofuels and bulk chemicals, including ethanol, higher alcohols, fatty acids, enzymes, amino acids, shikimate-derivatives, terpenoids, polyketides, and polymer precursors like 1,4-butanediol [8]. For higher productivity, metabolic engineering for the generation of these bio-chemicals necessitates substantial regulation of cellular metabolism. Due to tremendous advancements in biotechnology that have recent considerable advancements in biotechnology, which have various advantages, have accelerated the field of genetic engineering at a never-before-seen rate. Genetic and biological research have undergone a revolution as a result of the unique ability to precisely manipulate and alter the genomes of living things and the acceleration of the study of functional genomics [9]. In order to undertake time-saving sequential or multiplex alterations during genome editing, effective tools such as CRISPR-CAS system are required. The clustered regularly interspaced short palindromic repeats (CRISPR)/CRISPR-associated (Cas) system has recently been employed as a powerful genome engineering tool in a variety of prokaryotes and eukaryotes, including but not limited to *E. coli* [10]. There are numerous *E. coli* gene-editing methods available, but each has distinct advantages and disadvantages. Through the use of the Cas9 nuclease enzyme from *Streptococcus pyogenes*, targeted DNA restriction and DNA repair are achieved throughout this genetic engineering procedure. A blunt double-strand break (DSB) is created at a particular target locus when molecules of guide RNA bind Cas9 and the target sequence [11]. The CRISPR-Cas9 method has been shown to apply allelic exchange in *E. coli* with an efficacy of up to 65% [12], and to regulate gene expression by using a Cas9 protein lacking in nuclease [13]. Due to its many benefits, including its affordability, simplicity, high efficiency, and speed, CRISPR-Cas9 technologies have surpassed other earlier techniques like TALENs and ZFNs. The alteration of *E. coli* to make 1,3-propanediol, which was created by Genencor and DuPont and resulted in a commercial method, is a crucial illustration of a successful metabolic engineering effort [14]. Therefore, in this study, an effort was made to characterize the production of lipase from a Crispr-Cas9 edited *E. coli* and unedited *E. coli* with respect to the glucose source, media and different expression vectors in shake flask experiment.

Potato peel waste disposal is a critical issue due to its influence on the environment, such as decomposing organic waste in landfills and release methane, a powerful greenhouse gas that accelerates climate change, and as well potential loss of resources because they have multiple uses and are nutrient-rich, so disposing them away as waste is a loss of potential value and economic loss. By decreasing food waste and encouraging sustainable consumption and production, repurposing this waste material for industrial use, more especially, for the fermentation-based production of enzymes addresses the Sustainable Development Goals (SDG 2 and SDG 12). Effective lipase enzyme production at an industrial scale is necessary due to its several uses, including the manufacturing of biodiesel, food processing, and pharmaceuticals.

However, a number of obstacles prevent the best lipase production, including but not limited to low or poor yield and inconsistent quality, therefore a need arises to provide a promising method in lipase manufacturing in order to meet the expanding demand for lipase enzymes across a variety of industries.

## 2.0 Materials and Method

Out of the Blue CRISPR Kit with catalog number12012608EDU was purchased from Bio-Rad Research Company, United State of America. Potato peels waste were gathered from a nearby potato chip business at Akure, Ondo State, Nigeria. All other reagents used were obtained commercially and of analytical grade.

### 2.1 CRISPR-Cas9 Genome Editing of *Escherichia coli*

#### 2.1.1 LB Agar plate preparation

A 500 mL Kanamycin, IPTG (Isopropyl beta -D-1-thiogalactopyranoside) and X-gal (5-bromo-4-chloroindol-3-yl-b-D-galactopyranoside) (KIX) flask and a 1 Liter Kanamycin, IPTG, X-Gal/Spectinomycin (KIX/SPT) flask all labeled. In a vial, arabinose was completely dissolved by adding 3.0 ml of distilled water and stirring. 500 μl of distilled water was pipetted into another vial to dissolve the streptomycin. After giving the vial a thorough rinse, the KIX Mix slurry was transferred into the 700 ml of distilled water-labeled flask marked KIX/SPT. The insoluble powder was suspended by swirling the solution before a portion was transferred to the 500 ml KIX flask. Heat was applied to the LB agar powder until it boiled, with 7g placed in the KIX flask and the remaining amount in the KIX/SPT flask. IX, IX/ARA (arabinose) and IX/SPT plates were made with the corresponding mixtures. The stacks of plate were wrapped in aluminum foil and stored in an inverted position inside the refrigerator at 4^°^C. This was done seven days before the gene editing activity.

#### 2.1.2 Rehydration of bacteria

50 ml of distilled water and one LB broth capsule were placed inside a 150–250 ml bottle. This bottle was autoclaved three times with its cap loose in order to bring it to a boil. After then, it was allowed to reach 25^°^C. Using a sterile pipette tip, 250 milliliters from this solution were added to the vial containing the preserved *E. coli* HB101-pBRKan. To resuspend the organism, *E. coli*, the vial was lightly stirred and cultured at 37°C for 24 hours. Two days before the *E. coli* gene editing activity, this was accomplished.

#### 2.1.3 Streaking and incubation of starter plates

Resuspended *E. coli* was inoculated into eight IX and IX/ARA plates using a sterile plastic inoculation loop. The first streak pattern was diagonally oriented near the quadrant’s perimeter. After rotating the plate by one-fourth turn, the loop was moved from one side to another across the previously formed patterns several times before moving on to the next sector. There were two additional rounds of this treatment. After being cultured in an upside-down orientation at 37°C for 24 hours in an incubator oven, the starter plates were kept at 4°C. This was achieved 24 hours before the gene editing activity.

#### 2.1.4 Preparation of Plasmids

25 μl of pLZDonor plasmid was introduced to 8 PD tubes and 25 μl pLZDonor Guide plasmid to every PDG tubes. All solutions were stored in a refrigerator at 4°C until they were needed. All the reagents, with the exception of the transformation solution (TS), were brought to room temperature before use.

#### 2.1.5 Gene editing of *lacZ* gene in *E. coli* HB101

The icy transformation solution containing 250 μl was introduced to each of the four labeled microcentrifuge tubes (A–D) with a label. Bacterial colonies were individually dispersed from IX and IX/ARA plates into tubes A, B, C, and D. Tubes A and C received 10 μl of the pLZDonor (PD) plasmid, while tubes B and D received 10 μl of the pLZDonorGuide (PDG) plasmid. After 10 minutes on ice, the tubes experienced heat shock for 50 seconds at 60°C, and then they were returned to the ice. 250 μl of LB nutritional broth were added to each tube, which was then allowed to sit at room temperature for 20 minutes. IX/SPT plates A–D were equally covered with samples from tubes A–D, respectively. The plates were then incubated for 24 hours at 37°C to promote bacterial growth. The blue-white screening technique was used to confirm the success of *lacZ* editing.

#### 2.1.6 Genomic DNA extraction from bacteria

Five screw-cap tubes were labeled as SPE, UE, ED1, ED2, and ED3. After the beads were evenly resuspended by inverting the Insta-Gene Matrix, 250 μl was pipetted into each tube. Individual colonies were selected from designated plates, a blue colony from the IX/ARA plate agitated in tubes S, another blue colony from plate C into tube UE. Furthermore, individual white colonies from plate D were swirled in tubes ED1, ED2, and ED3. The tube caps were vortexed or flicked to mix after making sure they were closed. The tubes were then placed in a water bath and incubated for predetermined times and temperatures. This was followed by a cooling phase and more mixing.

#### 2.1.7 Design of donor DNA template and guide RNA

##### a) Guide RNA

The Cas9 endonuclease enzyme uses sgRNA to locate the target DNA. The guidance region of the Cas9 enzyme is designed to be complementary to the target DNA sequence in order to tell it where the cutting sites are.

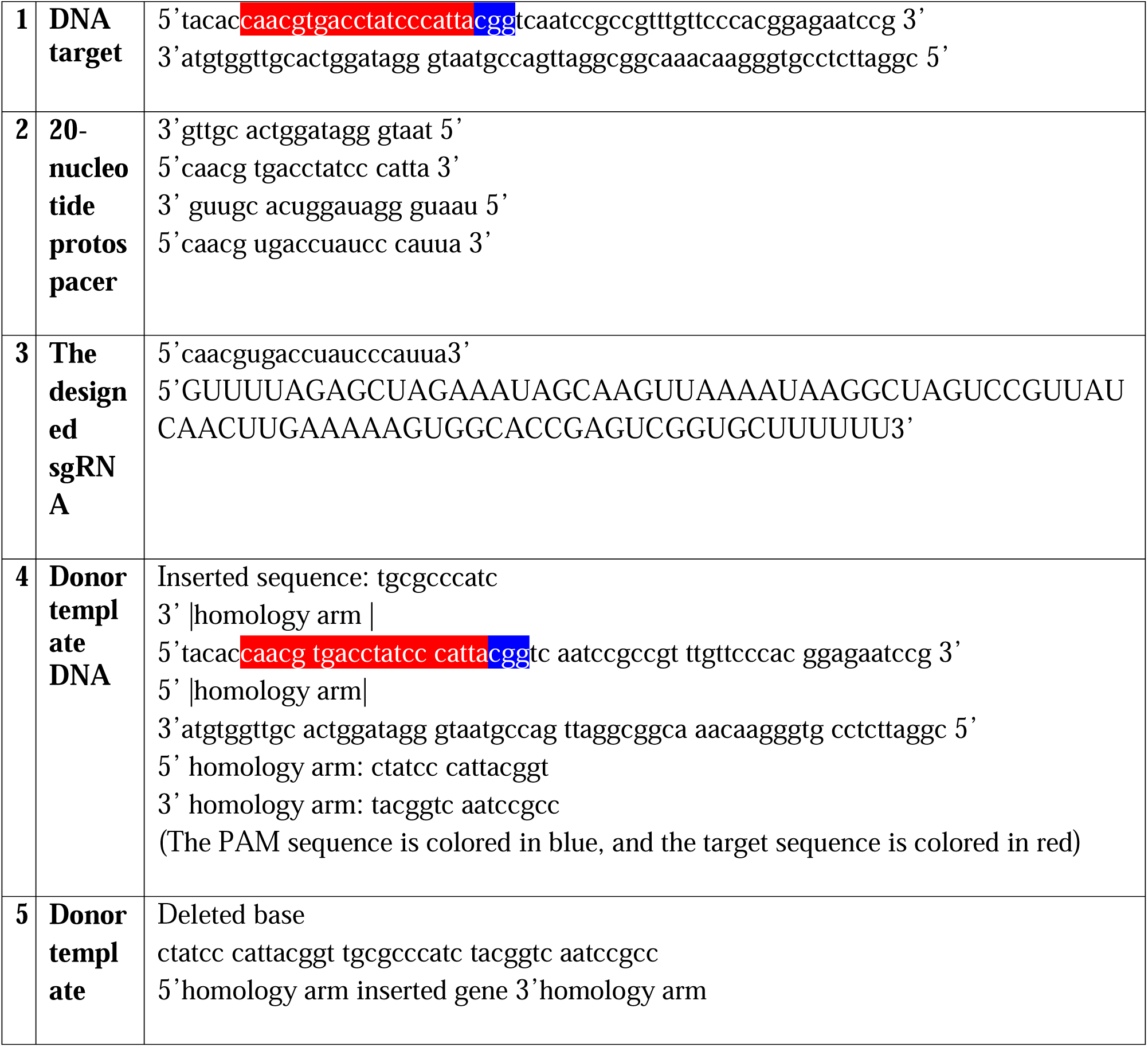

#### 2.1.8 Polymerase Chain Reaction

PCR was done to confirm DNA modifications. The tubes were centrifuged, and 10 μl of mastermix with primers (MMP) was aliquot to 7 PCR tubes that had been tagged Starter plate *E. coli* (SPE), Unedited *E. coli* (UE), Edited *E. coli* 1 (ED1), Edited *E. coli* 2 (ED2), Edited *E. coli* 3 (ED3), Negative control (-) and Positive control (+). Positive alongside negative PCR controls were pipetted into separate PCR tubes along with 10 μl of the screw-cap tubes supernatant. Ultimately, the thermal cycler was filled with the capped tubes and set to operate in accordance with initial denaturation of 94°C for 5mins then after 35 amplification cycles, denaturation of 94°C for 30secs, annealing temperature set at 62°C for 30secs followed by extension at 74°C for 5mins.

Multiplex PCR was used to test the samples for the desired DNA sequence presence. In this experiment, multiple amplicons are amplified simultaneously in a single reaction using a different set of primers for each amplicon. The majority of PCR tubes yielded a single blue colony from either the samples of product from the prior phase or the IX/ARA plate. From the latter source, the rest also received a white colony apiece.

#### 2.1.9 Gel Electrophoresis

The PCR amplification product that was produced was subjected to separation using a 1.0% agarose gel that also included ethidium bromide. In the solidified agarose gel that was immersed in 0.5X tris-borate-EDTA buffer in the electrophoresis tank, the PCR product was combined with a loading dye and then gently loaded onto wells. The electrophoresis was then conducted for 30 minutes at 100V. Following that, under UV light, the separated bands were seen in relation to the DNA ladder, which served as a molecular weight marker.

### 2.2 Enzyme production

#### 2.2.1 Substrate Preparation

The potato peels utilized in this study were acquired by means of manual peeling of Irish potatoes that were purchased from Shasha Market in Akure, Ondo State, Nigeria. The potatoes were cleaned with water and then rinsed with distilled water prior to peeling. In addition, the peels were cleaned in sterile distilled water and allowed to dry for 4 hours at 105°C in the oven. The dehydrated peels were ground in the laboratory, sieved to produce flour with a 0.5 mm mesh size, and stored in preparation for submerge fermentation [15].

#### 2.2.2 Lipolytic Efficiency of *Escherichia coli* on Lipase medium

The CRISPR Cas9 edited *E. coli* and unedited *E. coli* were separately inoculated on lipase producing media. The two cases’ medium were kept at pH 7.0, and five days of incubation were spent at 37°C. Any clear area that formed around the bacterial colony was an indication that lipolytic activity, or extracellular lipase, was being produced. The lipase medium was made as follows; KH_2_PO_4_ (1g/L), CaCl_2_ (2g/L), K_2_HP0_4_ (0.3g/L), (NH_4_)SO_4_ (2g/L), NaCl (5g/L), Olive Oil (1m/L), MgSO_4_ (0.2g/L), Methyl Red (0.0025g/L), Peptone (10g/L), Yeast Extract (5g/L) [16].

#### 2.2.3 Preparation of Potato Peel medium

A potato peel medium was made using the milled potato peel as the glucose source. This was done by preparing KH2PO4 (2.13g/L), NaHPO4 (1.5g/L), MgSO4 (0.5g/L) and the potato peel (1g/L). The medium was then adjusted to suitable pH and sterilized by autoclaving in 100ml conical flasks each having 75ml of the medium.

#### 2.2.4 Submerged Fermentation

After adjusting the pH of the three potato peel media to 7.0, they were incubated at three different temperatures: 10°C, 30°C, and 45°C. The medium in the various flasks were then inoculated with CRISPR Cas9 edited *E. coli* and unedited *E. coli* and subjected to various conditions listed above for 15 days with optical density (O.D) values taken every 3 days. This was done to observe the formation of cell mass as well as the mineralization of the potato peel in the medium which is acting as the glucose source. The contents of the flask were filtered using Whatman No. 1 filter paper and centrifuged for 20 minutes at 4,000 rpm on the filtrate. Crude enzymes were separated from supernatants and stored at 4°C for the enzyme assay [17].

#### 2.2.5 Partial Purification of Lipase

The protein solution was heated to 4°C and then put into a beaker fitted with a magnetic bar after submerged fermentation. A milliliter of the protein solution was taken, and 0.6g of ammonium sulphate was added. The protein solution was then agitated, and before adding the next portion, a small amount of ammonium sulphate was added and allowed to dissolve. At last, the beaker was left to stand for the entire night [17].

#### 2.2.6 Characterization of the Enzyme

##### 2.2.6.1 Temperature Effect on the Activity of Enzymes

Using Guaiacol (10 mM) as the substrate dissolved in sodium acetate buffer (10 mM pH 5.0) and incubated at 25^°^C, 35^°^C, 45^°^C, 55^°^C, and 65^°^C, the absorbance of the enzyme-catalyzed reaction was recorded to assess the impact of temperature on lipase activity. The reaction mixture was incubated for 15mins. The temperature at which the enzyme showed maximum activity was noted as the optimum temperature of the enzyme [18].

##### 2.2.6.2 pH Effect on the Activity of Enzymes

Using Guaiacol (10 mM) as the substrate, which was dissolved in buffers with varying pH values (acetate buffer, pH 4, pH 5, phosphate buffer, pH 6, pH 7, and tris-HCl buffer, pH 8) and incubated at 25°C for 15 minutes, the absorbance of an enzyme-catalyzed reaction at optimal temperature was recorded to examine the impact of pH on lipase activity. At 340 nm, absorbance was measured. [18].

#### 2.2.7 Statistical Analysis

One-way analysis of variance (ANOVA) was used to compare the enzymatic activity of the lipase enzyme made from CRISPR Cas9 edited and unedited *E. coli*. GraphPad Prism (version 9.3.1) was used to analyze the data, and Excel 2016 was used to visualize the results. P<0.05 served as a threshold for statistical significance.

## 3.0 RESULTS

CRISPR Cas9 gene editing of *lacZ* gene in *E. coli* was carried in four different petri dish. The observations of the petri dish after 24 hours of incubation at 37^°^C are shown in plate 1–4.

The Petri dish made up with containing colonies of unedited *E. coli* subjected to CRISPR-Cas9 system with Cas9 but without the single guide RNA (sgRNA) and arabinose, the repair machinery is shown in Plate 1a. A unique blue phenotype is displayed by the colonies; counting the colonies that have been observed, 116 colonies have this distinctive blue pigmentation.

Plate 1b presented the petri dish of CRISPR-Cas9 edited but not repaired *E. coli,* with sgRNA, Cas9 but without arabinose, the repair machinery, this was conspicuously devoid of observable colonies. Upon assessment, the count of blue or white colonies of *E. coli* on this plate was nil, registering a total count of zero. Plate 1c shows an illustration of unedited *E. coli* colonies subjected to the CRISPR-Cas9 system with Cas9, arabinose, the repair machinery but without sgRNA. These colonies exhibit a phenotype that is distinctly blue.

A total of 239 colonies were found to have this particular blue pigmentation after observation. The petri dish with CRISPR-Cas9 edited *E. coli* containing Cas9, sgRNA and arabinose the repair machinery on Plate 1d showed colonies with a characteristic white phenotype. 79 white *E. coli* colonies in all were found and counted after the enumeration.

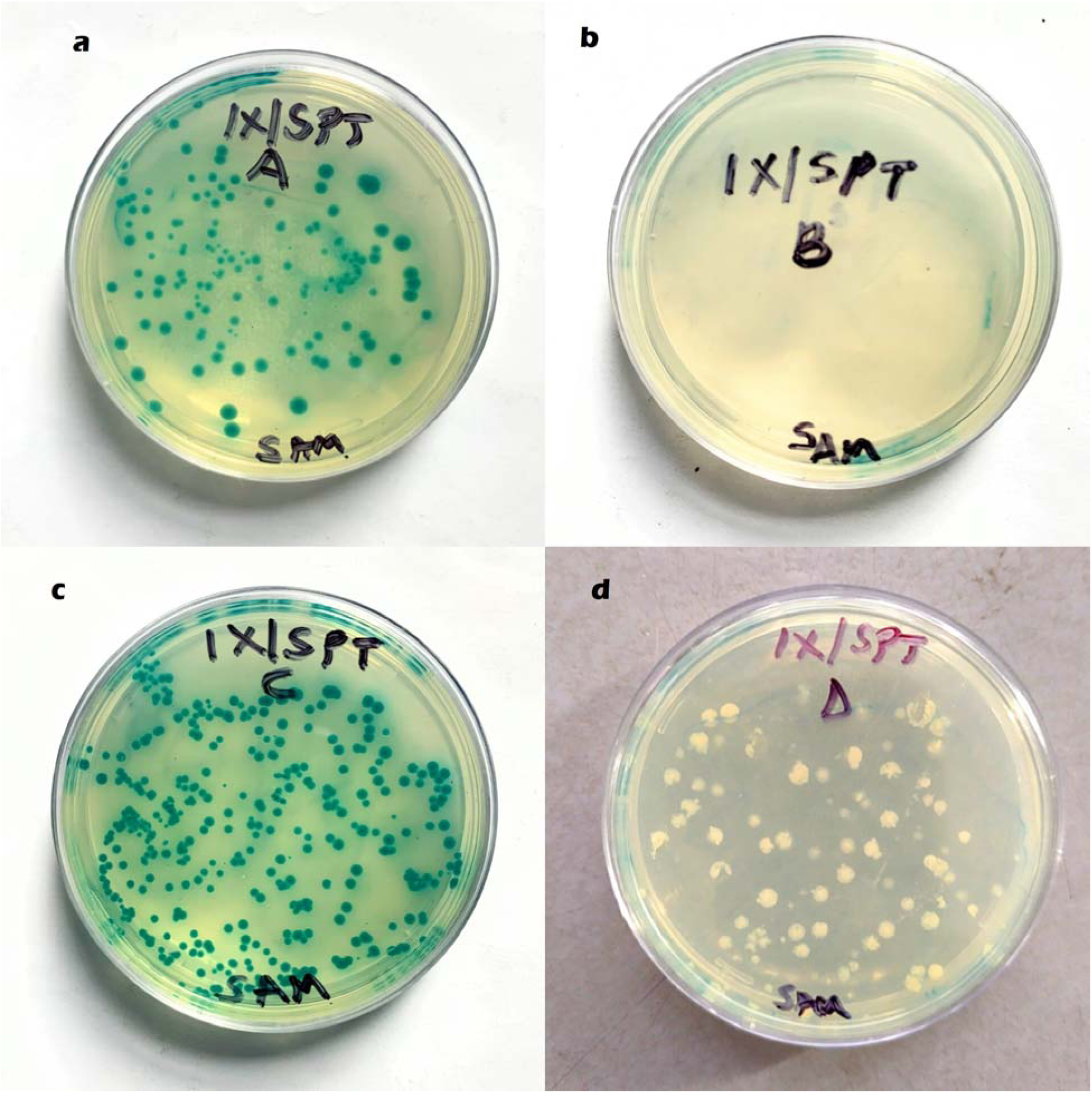

**Plate 1a:** Petri dish showing CRISPR Cas9 unedited *E. coli* with Cas9 enzyme but without single guide RNA (sgRNA) and repair machinery (arabinose) exhibiting blue colonies.

**Plate 1b:** Petri dish showing CRISPR Cas9 edited with Cas9 enzyme and sgRNA but not subjected to repair machinery (no arabinose) with no colony of *E. coli*.

**Plate 1c**: Petri dish showing CRISPR Cas9 unedited *E. coli* with Cas9 enzyme and active repair machinery (arabinose) but without single guide RNA (sgRNA) exhibiting blue colonies.

**Plate 1d:** Petri dish showing CRISPR Cas9 edited *E. coli* with Cas9 enzyme and sgRNA subjected to active repair machinery showing white colonies.

Plate 2 illustrates the electrophoretic band pattern obtained from the CRISPR-Cas9 experiment. Within lanes Edited *E. coli* 3 (ED3), Edited *E. coli* 2 (ED2), and Edited *E. coli* 1 (ED1), originating from the *E. coli* displaying a white phenotype. Two distinct bands were observed with band size of 650 bp. Two bands, originating from *E. coli* with a blue phenotype, were also detected in starter plate *E. coli* (SPE) and unedited *E. coli* (UE) lanes with band size of 1,100 bp. In the positive control, three distinct bands were observed with 1,100 bp, 650 bp and 350 bp all present, whereas in the negative control, no bands were detected.

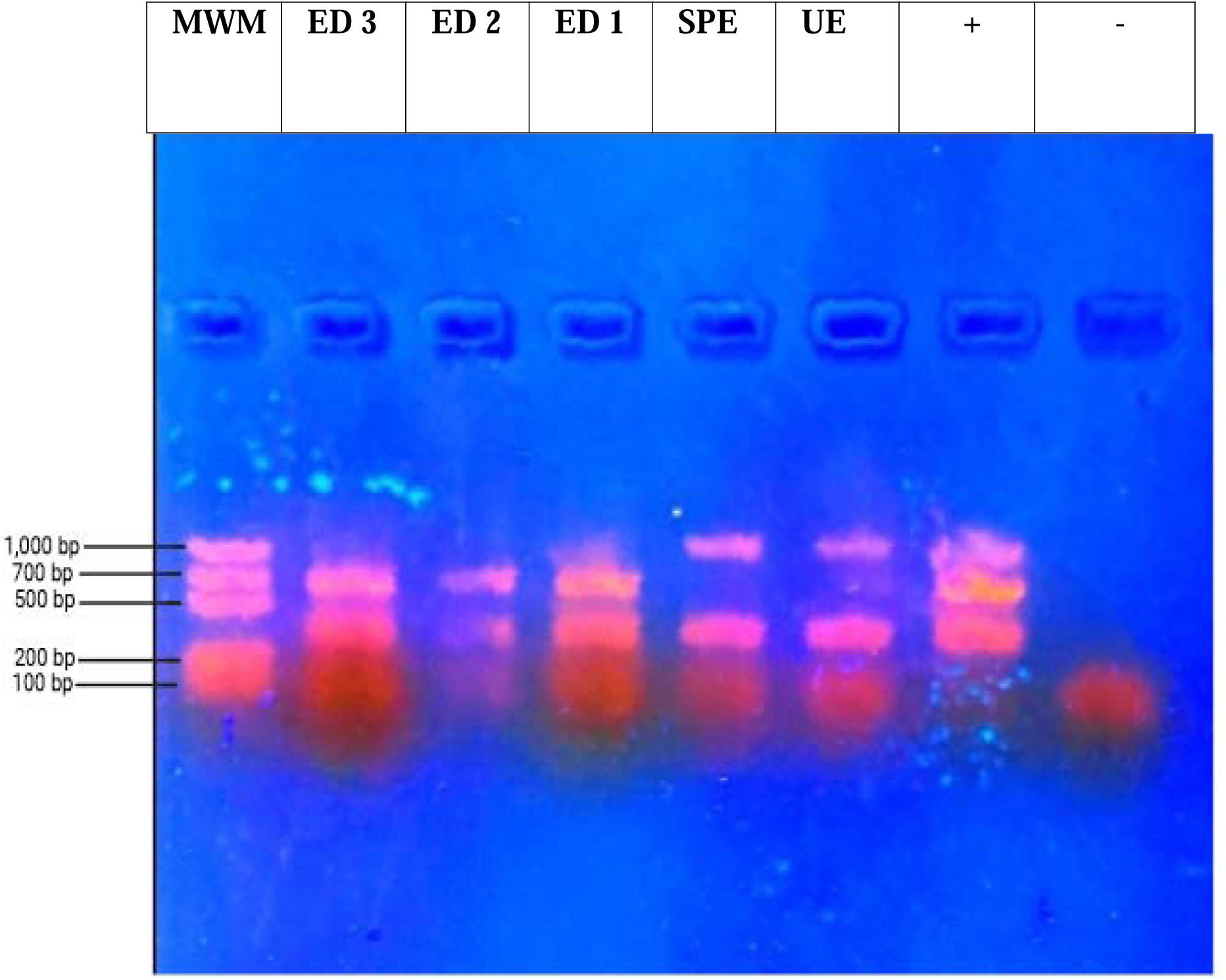

**Plate 2:** Electrophoretic bands of the CRSIPR Cas9 edited *E. coli* and unedited *E. coli* PCR products.

**KEY**

MWM= Molecular weight marker.

ED3=Edited *E. coli* 3, ED2=Edited *E. coli* 2, ED1=Edited *E. coli* 1.

SPE= Starter plate *E. coli*, UE= Unedited *E. coli*.

+ = Positive control, - = Negative control.

Following five days of incubation, lipase media containing both CRISPR Cas9-edited and unedited *E. coli* were inspected for a clear zone to determine whether lipolytic activity was present. Plate 3 shows the CRISPR Cas9 edited *E. coli* petri dish had a clear zone surrounding the colonies in the lipase medium observed more than the CRISPR unedited *E. coli*. Plate 4a shows crude enzyme extract obtained after submerged fermentation of potato peel with CRISPR Cas9 edited *E. coli* and unedited *E. coli*. The lipase enzyme extracted from CRISPR Cas9 edited and unedited *E. coli* was partially purified using ammonium sulphate. Plate 4b showed the purified lipase enzyme retrieved from both varieties of *E. coli*.

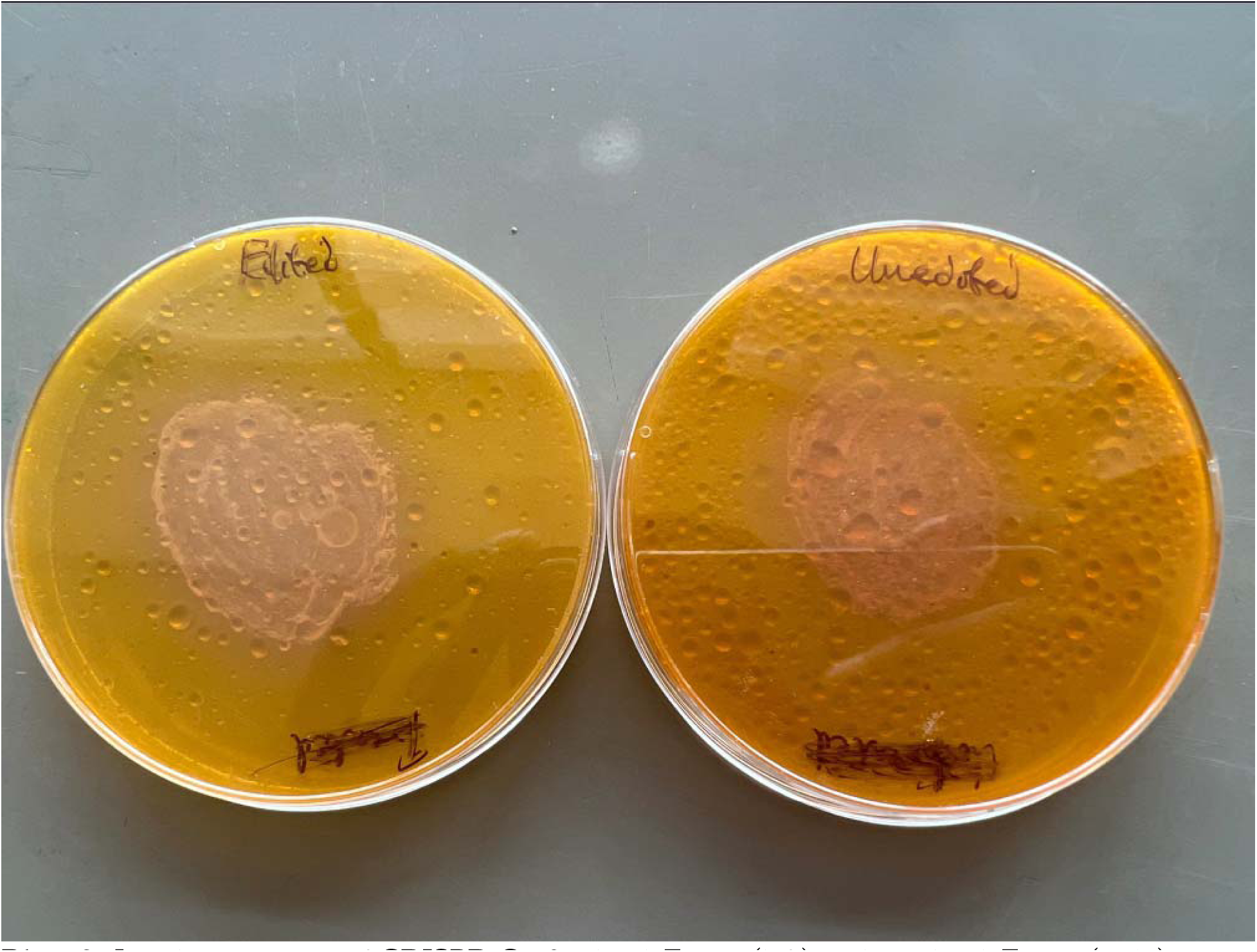

**Plate 3:** Lipolytic activity of CRISPR Cas9 edited *E. coli* (left) and unedited *E. coli* (right) on lipase medium.

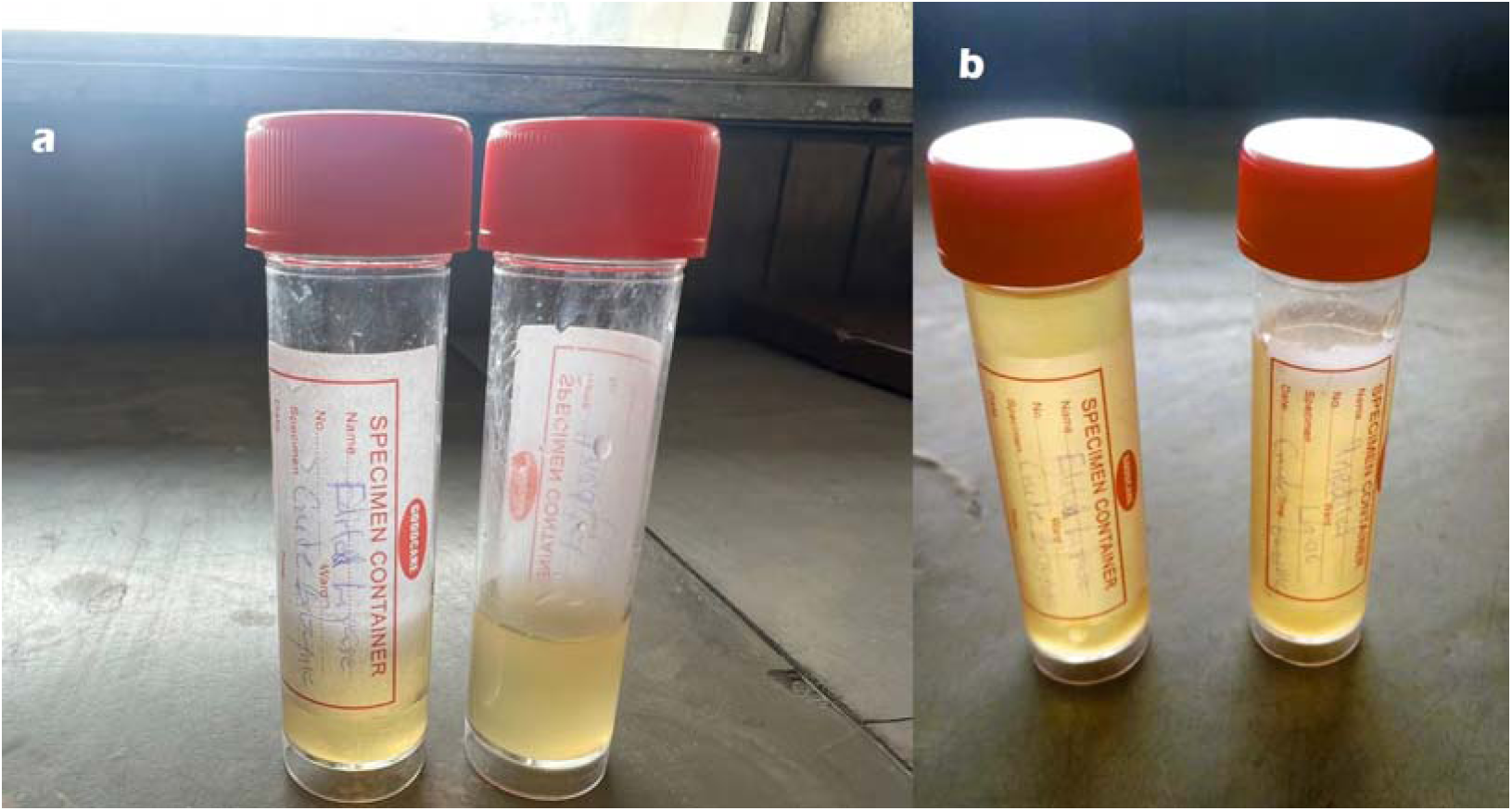

**Plate 4a:** Crude enzyme extract after submerged fermentation with CRISPR Cas9 edited *E. coli* and unedited *E. coli*.

**Plate 4b:** CRISPR Cas9 edited *E. coli* and unedited *E. coli* enzyme extract after partial purification using ammonium sulphate.

The growth of CRISPR Cas9 unedited *E. coli* in relation to temperature in the potato peel medium under 15 days of incubation at 10°C, 30°C, and 45°C was presented in Figure 1 showing an increased cell mass at 10°C. Figure 2 shows the CRISPR Cas9 edited *E. coli* exhibited an increased cell mass concentration at 10°C among the different temperatures. Figure 3 illustrates the impact of temperature on the pure lipase derived from potato peels by fermentation with both CRISPR Cas9 modified and unedited *E. coli*. As the temperature increased from 25^°^C to 35^°^C, the activity of the lipase produced by the CRISPR Cas9 unedited and edited *Escherichia coli* increased significantly (p<0.05). Significant (p>0.05) decreases in enzyme activity were observed in both purified lipase enzymes produced by CRISPR Cas9 edited and unedited from 45^°^C to 65^°^C with further temperature increase. The lipase enzyme synthesized from both CRISPR Cas9-edited and unedited *E. coli* showed peak activity at 35°C and lowest activity at 65°C. At a 95% confidence interval, the difference in temperature’s influence on the purified lipase derived from potato peels through fermentation using CRISPR Cas9 edited *E. coli* and unedited *E. coli* is statistically significant (p<0.05). The effect of pH on the purified lipase enzyme obtained from potato peels through fermentation with both CRISPR Cas9 edited and unedited *E. coli* was demonstrated in Figure 4. From pH 7 to pH 8, a substantial (p<0.05) decrease in activity was seen. The purified lipase enzyme generated with CRISPR Cas9 edited *E. coli* and unedited *E. coli* was found to exhibit maximum enzyme activity at pH 7 and minimum enzyme activity at pH 4. The difference in effect of pH on the purified lipase obtained from potato peels by fermentation with CRISPR Cas9 edited *E. coli* and unedited *E. coli* is significant (p<0.05).

**Figure 1:**
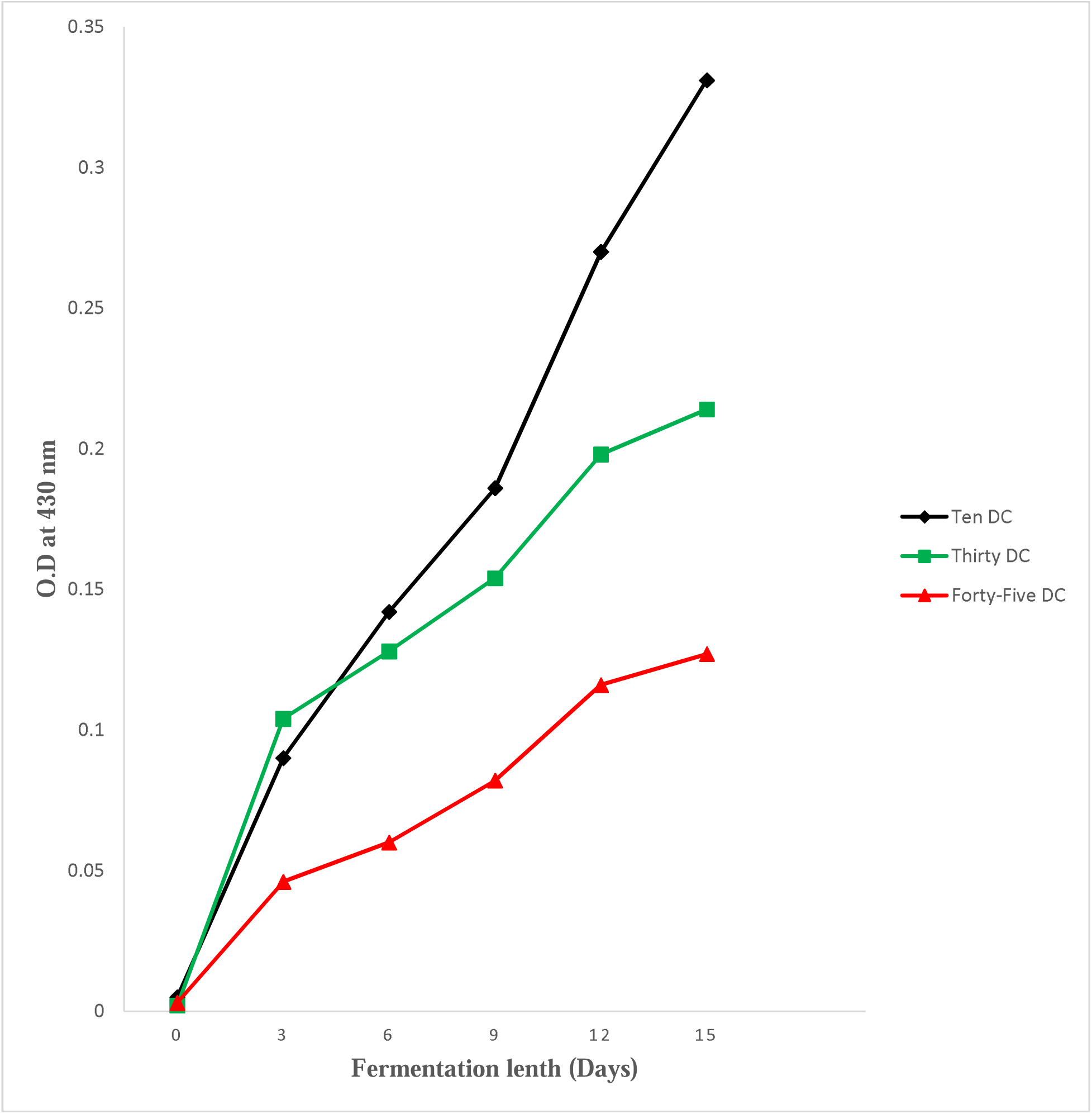
Effect of temperature on CRISPR Cas9 unedited *Escherichia coli* in potato peels medium. (Every value plotted denotes the average of two measurements ± SD). Key-Ten DC: 10^°^C, Thirty DC: 30^°^C, Forty-Five DC: 45^°^C.

**Figure 2:**
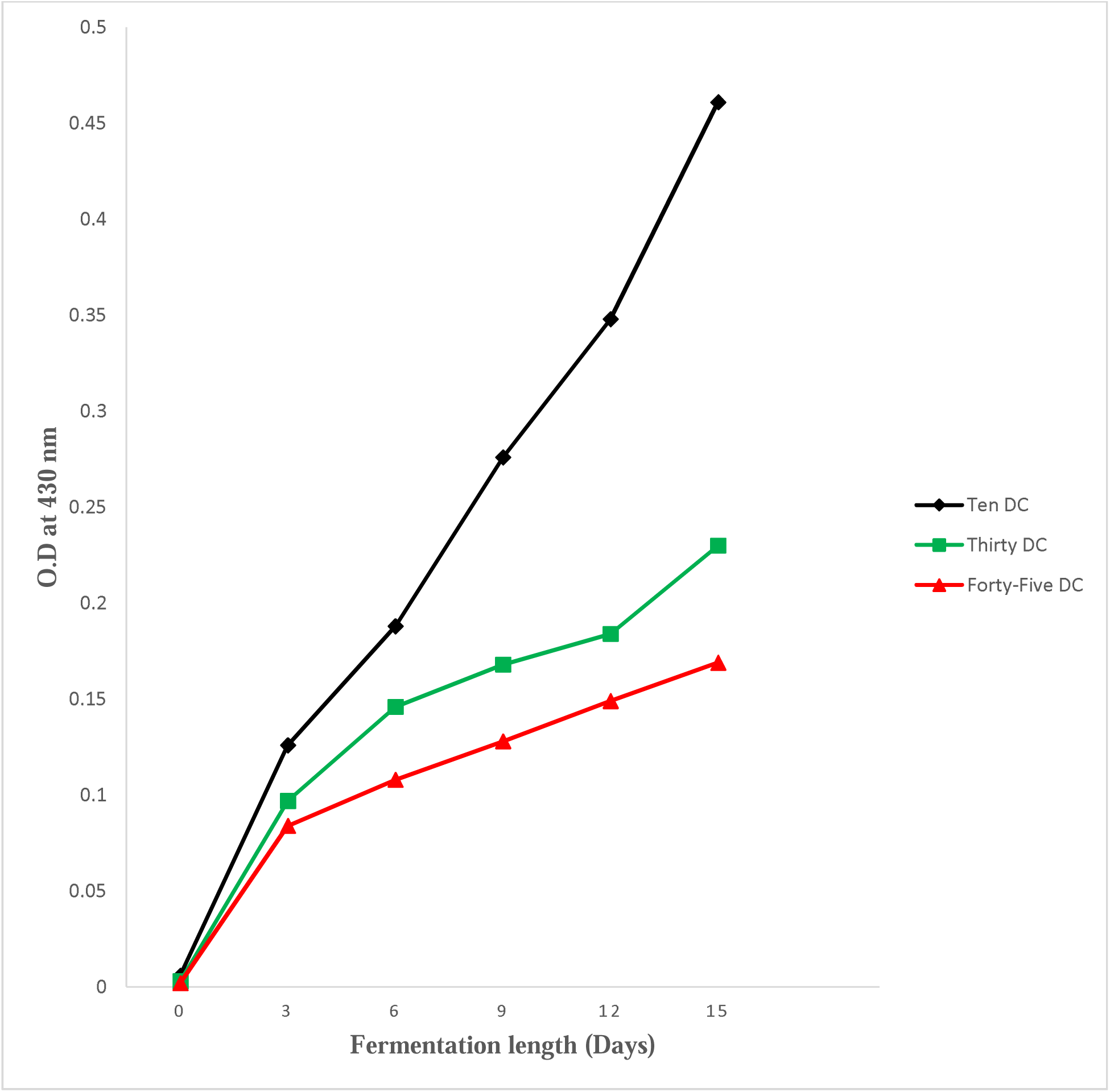
Effect of temperature on CRISPR Cas9 edited *Escherichia coli* in potato peels medium. (Every value plotted denotes the average of two measurements ± SD). Key-Ten DC: 10^°^C, Thirty DC: 30^°^C, Forty-Five DC: 45^°^C.

**Figure 3:**
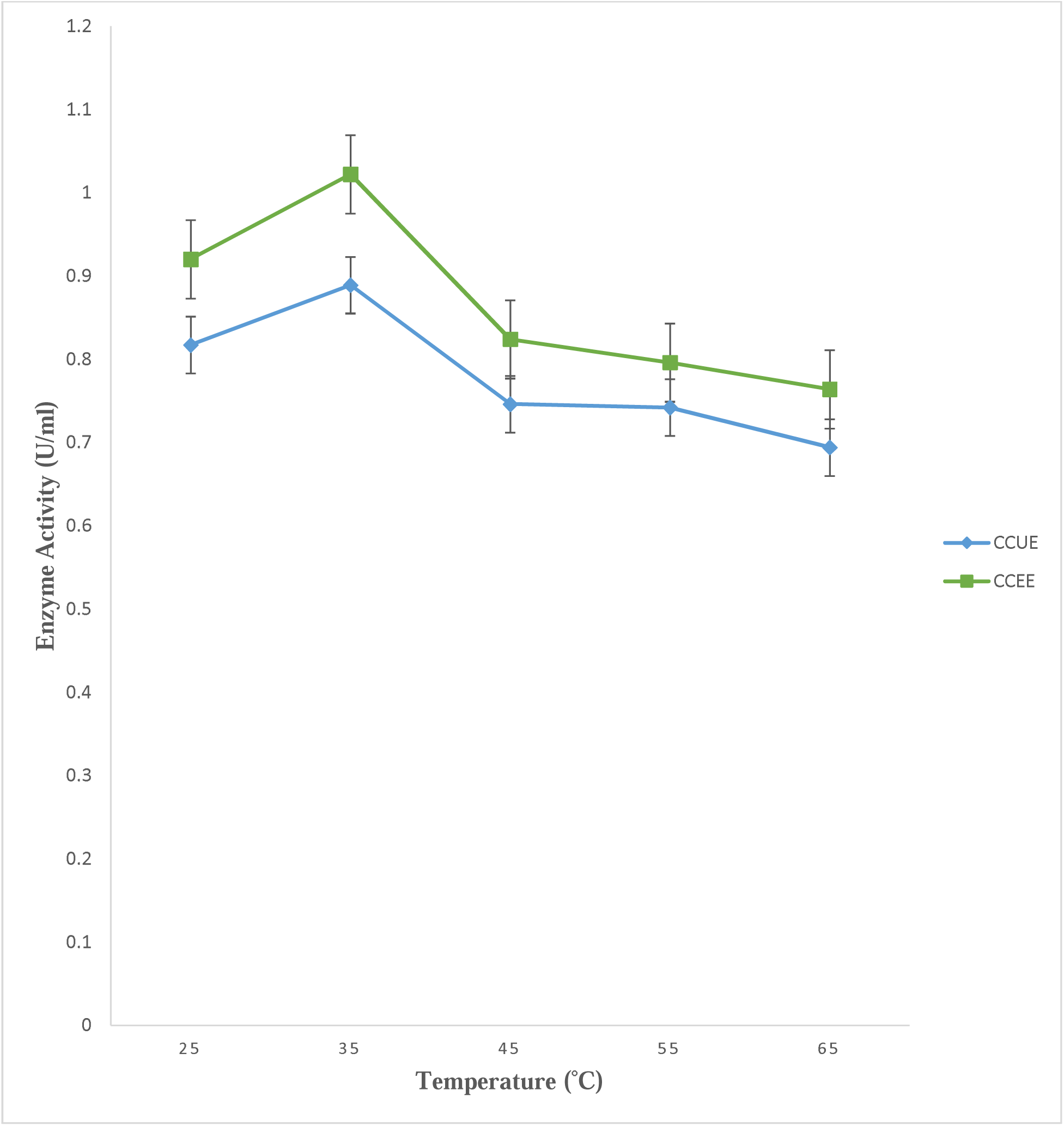
Temperature effect on the activity of enzymes on purified lipase enzyme extracted from potato peels with CRISPR Cas9 edited *E. coli* and unedited *E. coli*. (Every value plotted denotes the average of two measurements ± SD). Key: CCUE: CRISPR Cas9 Unedited *Escherichia coli*, CCEE: CRISPR Cas9 Edited *Escherichia coli*.

**Figure 4:**
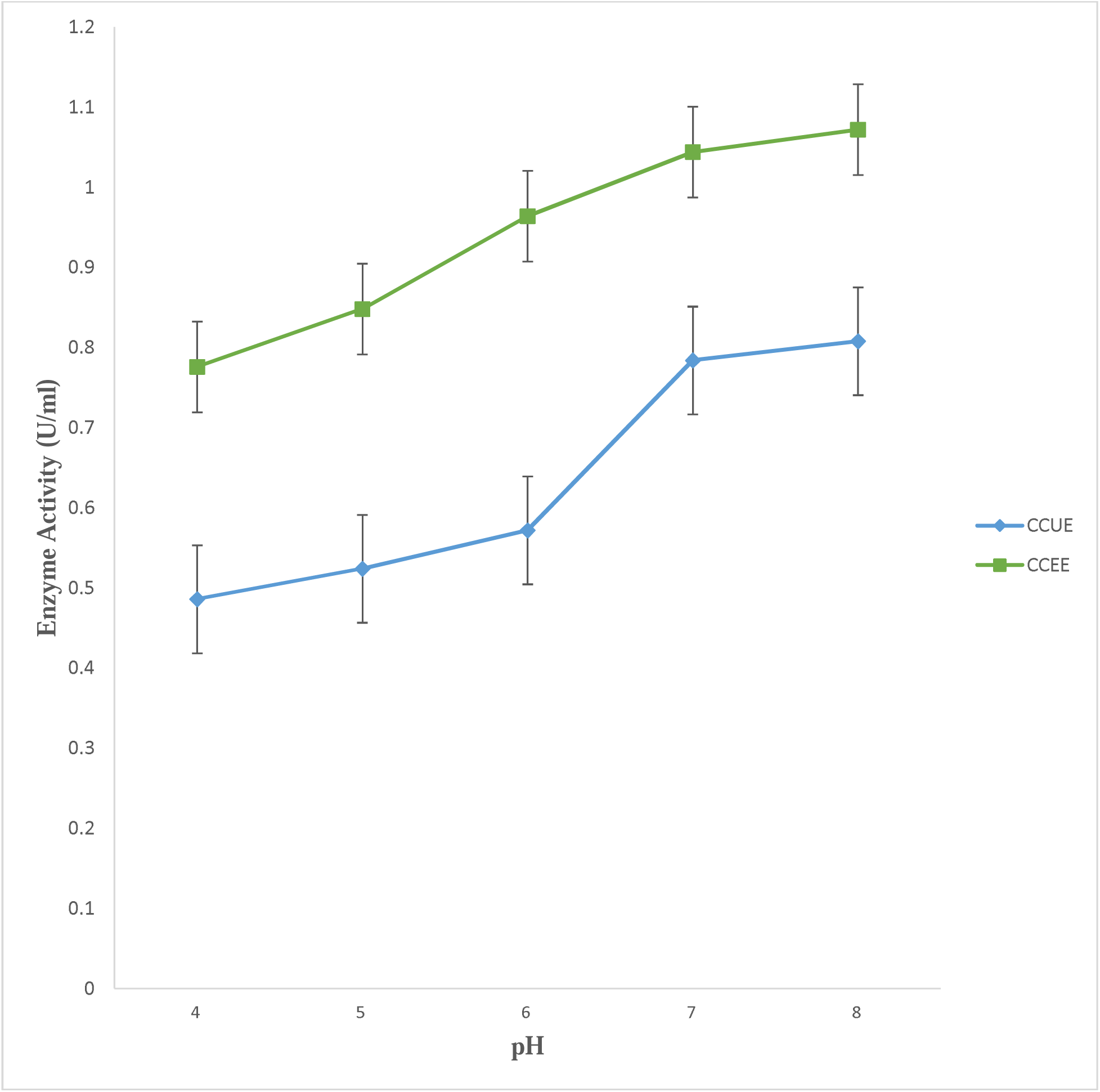
pHs effect on the activity of enzymes on purified lipase enzyme extracted from potato peels with CRISPR Cas9 edited *E. coli* and unedited *E. coli*. (Every value plotted denotes the average of two measurements ± SD). Key: CCUE: CRISPR Cas9 Unedited *Escherichia coli*, CCEE: CRISPR Cas9 Edited *Escherichia coli*.

## 4.0 DISCUSSION

These days, the more straightforward and accurate method of gene editing made possible by the latest advances in biotechnology that have led to the discovery of the CRISPR/Cas9 combination is preferred over other past genome editing technology [19]. Through the use of CRISPR-Cas9, *E. coli* strains can have specific genetic modifications made to them, thereby strategically improving the biosynthesis pathways involved in the production of enzyme, specifically lipase.

Using the CRISPR-Cas9 technology, the *lacZ* gene, which is normally present in the *E. coli* HB101-pBRKan chromosome, was altered in the first experiment. The results of the *lacZ* gene alteration show that when the bacteria were cultured on X-gal (5-bromo-4-chloroindol-3-yl-b-D-galactopyranoside)-containing media, both edited and unedited *lacZ* genes were present.

Bacteria on Plate A which was originated from a starter plate made up of kanamycin, IPTG (Isopropyl beta -D-1-thiogalactopyranoside) and X-gal but devoid of arabinose which is the repair machinery, and the *E. coli* containing the plasmid donor (DNA donor template), Cas9 but lack the single guide RNA (sgRNA) evade the Cas9 cleavage thereby having blue colony present which indicated that *lacZ* gene has not been edited. This blue colony coloration suggests the hydrolysis of X-gal by active beta galactosidase, which is only going to be expressed in the event that the *lacZ* gene is active and stimulated by IPTG. This result is in agreement with [10] findings who reported that the sgRNA required to guide Cas9 to cleave *lacZ* gene was not present in the bacteria transformed with plasmid donor and as a result no gene editing occurred and the transformants remained blue regardless of whether the repair system was turned off or on.

The absence of colonies on Plate B made up of kanamycin, IPTG and X-gal but devoid of arabinose which is the repair machinery, and the *E. coli* containing the plasmid donor guide (DNA donor template and sgRNA) and Cas9 demonstrated that the *lacZ* gene was edited but not repaired. These bacteria express the Cas9 enzyme as they were given sgRNA to direct Cas9 to cut the *lacZ* gene. Nevertheless, the repair system was not triggered because they were cultivated on starter plates devoid of arabinose. As a result, the bacteria died without growing colonies and the cut was irreparable. These findings validate the study reported by [20] that the bacteria on this plate have no functional repair machinery and have not survived the double stranded break, therefore, the break could not be repaired and the bacteria died without forming colonies. The 239 colonies with the plasmid donor (DNA donor template), Cas9 but lack the single guide RNA found on plate C made up of kanamycin, IPTG and X-gal and arabinose which is the repair machinery, indicated that the *lacZ* gene fails to be cut by the Cas9 thereby having blue colony present. This blue colony pigment suggests the reaction between X-gal and active beta galactosidase, which is only going to be expressed in the event that the *lacZ* gene is functional and triggered by IPTG. The fact that the stop codon inserted by the donor DNA template prevents the development of functional b-galactosidase and prevents the bacteria on this plate from being cut and later repaired is supported by the observation made by [21] in their investigation. The bacteria containing plasmid donor guide (DNA donor template and sgRNA), Cas9 subjected to Cas9 cleavage, on Plate D were grown on an arabinose-containing starter plate with kanamycin, IPTG and X-gal after being given sgRNA to instruct Cas-9 to cut the *lacZ* gene. As a result, the bacteria’s white color indicates that the *lacZ* gene has been cut and then repaired. This report aligns with the study reported by [22] who further discussed that the double strand break that occurred was repaired with the DNA donor template through repair mechanism known as homology directed repair (HDR) and also that Since the lack of blue color indicates no enzyme activity, the presence of X-gal and IPTG indicates that the *lacZ* gene is not active.

By using amplicon amplification, more details were discovered on any alterations to the DNA. If the donor DNA template is absent or the primer binding regions are damaged, the target PCR sequence in multiplex PCR cannot be amplified. Due to the primer sets, each PCR sample produced distinct amplicons. The 1,100 bp amplicon that was generated indicated that *lacZ* gene is functional because the Cas9 cut site could not be modified, which at the molecular level confirms the results discussed above about the CRISPR Cas9 unedited *E. coli* colonies found on plate A and C having an active *lacZ* gene. The 650 bp amplicon that was obtained demonstrated that the donor DNA was used to repair the intended cut site which further confirms the result of the CRISPR Cas9 edited *E. coli* found on plate D, that the *lacZ* gene is no longer functional because it has been edited and repaired. The 350 bp amplicon’s presence revealed that genomic DNA was extracted and amplified successfully irrespective if *lacZ* gene was changed or not.

Based on the amplicons, the gel electrophoresis assisted in visualizing the PCR products as DNA bands and provided information on whether the *lacZ* gene was altered. The presence of the luminous color band signified the success of the PCR test. These findings validate studies reported by [10] and [22] who both revealed in their study that the first primer (1,100 bp) was created to detect *lacZ* gene that had not been changed as the primer will not bind if the desired cut site has been altered, the second primer (650 bp) set was created to detect *lacZ* gene that had been changed, and as a control, the third pair of primers (350 bp) amplified an unrelated region well downstream of the *lacZ* gene to ensure that the sample contained chromosomal DNA.

For the purpose of producing lipase by submerged culture, numerous studies have been conducted to identify the ideal culture and nutritional needs. After editing the *E. coli* using CRISPR Cas9, the edited and unedited strains are screened for lipolytic activity using olive oil as the lipid substrate to produce lipase. The development of a transparent zone around bacterial colonies is a sign that the lipid substrate (olive oil) in the medium has hydrolyzed or broken down [23]. When lipids (fats and oils) are hydrolyzed into their constituent fatty acids and glycerol by lipase enzymes secreted by microorganisms, the lipids can diffuse away from the bacterial colonies, forming a clear zone where lipolytic activity has taken place [24]. It is possible that the CRISPR Cas9 edited *E. coli* without the *lacZ* gene has higher lipolytic activity in breaking down the olive oil present in the medium into fatty acids and glycerol because a more noticeable clear zone has been observed around the CRISPR Cas9 edited *E. coli* colonies of the when compared to the unedited *E. coli* with a functional *lacZ* gene as discussed by [25]. Using the same screening technique, a total of seven lipolytic bacteria were isolated by [26]. These findings align with numerous research on microbial lipases that demonstrate elevated lipase production when olive oil is present among the various oils examined [27].

In contrast to the results of [28] and [29], who both stated that 37°C was the highest biomass concentration, the study’s findings showed that at 10°C, a higher biomass concentration was seen in 10°C, 30°C, and 45°C which are some of the different temperature conditions in which the potato peel medium was inoculated with both CRISPR edited *E. coli* and unedited *E. coli*. This finding is subjected to a future research to determine why this specific strain of *E. coli* HB101-pBRKan ferments better at low temperatures as opposed to the optimal temperature, and what factors led to the strain *E. coli* HB101-pBRKan showing its maximum biomass concentration at 10°C. Following submerged fermentation, the crude enzyme was purified with ammonium sulphate, yielding a clear solution. [18] employed a similar procedure to treat the crude enzyme assay from their study, obtaining opaque solution following ammonium sulphate treatment. This study selected ammonium sulphate due to its accessibility and comparatively low cost, as reported by [16].

One environmental factor that can influence the activity of enzymes is temperature. This study found that the lipase derived from both CRISPR Cas9 edited *E. coli* and unedited *E. coli* exhibited maximum activity at 35°C. The results of this study are consistent with findings of [30], who found that bacterial lipase activity increased slightly with temperature and peaked at 35°C. However, it is in contrast to the findings of [31] who reported that the highest lipase activity was recorded at 50°C from *Microbacterium sp*.

The most crucial factor for producing enzyme activity is the pH of the medium. Lipase production rate is dependent on the starting pH of the growth medium. The highest activity for the lipase enzyme in this study was obtained at pH 7. This result is correlated with the findings of [32], who found that *Staphylococcus* showed maximal lipase activity at pH 7. Nevertheless, However, this contradicts a study by [31] that found lipase from *Microbacterium sp* had its maximum activity at pH 8.5, and a study by Lee *et al*. (2001) that found lipases from *Bacillus thermoleovorans* ID1 had their maximum activity at pH 9.

In this study, the lipase derived from the CRISPR-Cas9 edited *Escherichia coli* without a functional *lacZ* gene is demonstrated to have higher enzyme (temperature and pH) activity compared to the CRISPR-Cas9 unedited *E. coli* with a functioning *lacZ* gene. β-galactosidase, an enzyme that breaks down lactose, is produced by the *lacZ gene* in *E. coli,* but due to the alterations in metabolism and regulatory pathways, editing the *lacZ* gene with the CRISPR Cas9 system may have a variety of effects on *E. coli* cellular physiology. The synthesis of lipase may be favored by knocking out the *lacZ* gene, which could divert metabolic resources away from lactose metabolism and β-galactosidase production toward other pathway and purposes. Absence of the *lacZ* gene may impact cellular regulatory networks. This could result in modifications to gene expression patterns and the activation of lipase production-related pathways or changes the way lipase genes or associated metabolic pathways are regulated. Future research, such as comparative gene expression analyses and comprehensive metabolic profiling, would be required to acquire a more profound comprehension of the fundamental mechanisms responsible for the detected variations in lipase activity between the two strains.

## CONCLUSION

This study ascertained that *lacZ* gene editing of *Escherichia coli* with CRISPR Cas9 system optimize the CRISPR Cas9 edited *Escherichia coli* strain to make use of this waste substrate in producing lipase the more. This demonstrates the application of genome editing techniques for sustainable and eco-friendly enzyme production utilizing agricultural by-products. However, more in-depth research should be done to further investigate the specific pathways, regulatory elements, and interactions involved in the augmented expression of lipase-related genes post-CRISPR editing, promoting the creation of more streamlined and large-scale methods for the effective production of lipase.

## Declarations

### Ethics approval and consent to participate

Not applicable

### Consent to publication

Not applicable

### Availability of Data and Materials

The data sets used and/or analyzed during this study are available from the corresponding author upon reasonable request.

### Code Availability

Not applicable

### Authors’ contribution

Joseph Bamidele MINARI conceived the project and defined the research methodology; Joseph Bamidele MINARI, Idowu Samuel DADA, Dhikrullah Oluwatope, ABDULAZEEZ Gift NWOSU performed the experiments: Joseph Bamidele MINARI, * Idowu Samuel DADA wrote the paper.

## Acknowledgment

We thank the University of Lagos Central Research Laboratory for their technical support.

## REFERENCES

1. Abebaw, G. (2020). Review on: Its potentials and application of potato peel (Waste). Journal of Aquaculture and Livestock Production, 1(1), 1–4.

2. Chang, M. Y., Chan, E. S., & Song, C. P. (2021). Biodiesel production catalysed by low-cost liquid enzyme Eversa® Transform 2.0: Effect of free fatty acid content on lipase methanol tolerance and kinetic model. Fuel, 283, 119266–119274.

3. Bornscheuer, U. T. (2018). The fourth wave of biocatalysis is approaching, *Philosophical Transactions of the Royal Society A:* Mathematical, Physical & Engineering Sciences, 376, 2110–2117.

4. Ha, J., Park, J. Y., Choi, Y., Chang, P. S., & Park, K. M. (2021). Comparative analysis of universal protein extraction methodologies for screening of lipase activity from agricultural products. Catalysts, 11(7), 816–827.

5. Javed, S., Azeem, F., Hussain, S., Rasul, I., Siddique, M. H., Riaz, M., Afzal, M., Kouser, A., & Nadeem, H. (2018). Bacterial lipases: A review on purification and characterization. Progress in Biophysics and Molecular Biology,132, 23–34.

6. Bano, K., Kuddus, M., Zaheer, M. R., Zia, Q., Khan, M. F., Gupta, A., & Aliev, G. (2017). Microbial enzymatic degradation of biodegradable plastics. Current Pharmaceutical Biotechnology, 18(5), 429–440.

7. Khurana, J., Pratibha, C., & Kaur, J. (2017). Studies on recombinant lipase production by *E. coli*: Effect of media and bacterial expression system optimization. Molecular Biology, 2 (1)00008.

8. Yang, D., Prabowo, C. P. S., Eun, H., Park, S. Y., Cho, I. J., & Jiao, S., et al. (2021). *Escherichia coli* as a platform microbial host for systems metabolic engineering. Essays in Biochemistry, 65(2), 225–246.

9. Tavakoli, K., Pour-Aboughadareh, A., Kianersi, F., Poczai, P., Etminan, A., & Shooshtari, L. (2021). Applications of CRISPR-Cas9 as an advanced genome editing system in life sciences. BioTech, 10(3), 14–31.

10. Naeem, M. A. (2022). CRISPR/Cas9 gene editing in bacteria: Leading to hematopoietic Stem cell editing in Pakistan. Journal of Haematology and Stem Cell Research, 2(1), 4–7.

11. 11. Loureiro, A., & da Silva, G. J. (2019). CRISPR-Cas: converting a bacterial defense mechanism into a state-of-the-art genetic manipulation tool. Antibiotics, 8(1), 18–43.

12. Jiang, W., Bikard, D., Cox, D., Zhang, F., & Marraffini, L. A. (2013). RNA guided editing of bacterial genomes using CRISPR-Cas systems. Nature Biotechnology, 31, 233–239.

13. Qi, L. S., Larson, M. H., Gilbert, L. A., Doudna, J. A., Weissman, J. S., Arkin, A. P., & Lim, W. A. (2013). Repurposing CRISPR as an RNA-guided platform for sequence-specific control of gene expression. Cell, 152(5), 1173–1183.

14. 14. Yim, H., Haselbeck, R., Niu, W., Pujol-Baxley, C., Burgard, A., Boldt, J., & Van Dien, S. (2011). Metabolic engineering of Escherichia coli for direct production of 1, 4-butanediol. Nature chemical biology, 7(7), 445–452.

15. Maxwell, O., Chinwuba, U., & Onyebuchukwu, M. (2019) Protein enrichment of potato peels using *Saccharomyces cerevisiae* via solid-state fermentation process. Advances in Chemical Engineering and Science, 9(1), 99–108.

16. Kareem, S. O., Adebayo, O. S., Balogun, S. A., Adeogun, A. I., & Akinde, S. B. (2017). Purification and characterization of lipase from *Aspergillus flavus* PW2961 using magnetic nanoparticles. Nigerian Journal of Biotechnology, 32, 77–82.

17. Minari, J. B., & Agho, E.E. (2018). Laccase extraction, purification and characterization from potato peels. Bells University Journal of Applied Sciences and Environment, 1(2), 1–6.

18. Agho, E. E., Minari, J. B., Emelumadu, E. N., Adeniyi, O. A., & Olajig, F. R. (2022). Characterization of laccase from the fungi *Fusarium* isolated from potato peels using carbon and nitrogen sources. Advances in Technology, 2(2), 183–197.

19. Gan, W. C., & Ling, A. P. K. (2022). CRISPR/Cas9 in plant biotechnology: Applications and challenges. Biotechnologia, 103(1), 81–93.

20. Cinti, R. (2021). Using CRISPR gene editing to modify the *lacZ* gene in *E. coli*. GMVS Journal of Molecular Biology, 30-41.

21. Chai, R., Zhang, Q., Wu, J., Shi, Z., Li, Y., Gao, Y., & Qiu, L. (2023). Single-Stranded DNA-binding proteins mediate DSB repair and effectively improve CRISPR/Cas9 genome editing in *Escherichia coli* and *Pseudomonas*. Microorganisms, 11(4), 850–865.

22. Vo, B. (2022). Winning entry for 2022. Library Research Prize. 18. https://digitalcommons.conncoll.edu/libprize/18.

23. Abubakar, A., Abioye, O. P., Aransiola, S. A., Raju, M. N., & Prasad, R. (2024). Crude oil biodegradation potential of lipase produced by *Bacillus subtilis* and *Pseudomonas aeruginosa* isolated from hydrocarbon contaminated soil. Environmental Chemistry and Ecotoxicology, 1-26.

24. Bharathi, D, C., & Rajalakshmi, G. (2019). Microbial lipases: An overview of screening, production and purification. Biocatalysis and Agricultural and Biotechnology, 22, 101368–101375.

25. Yan, J., Yan, Y., Madzak, C., & Han, B. (2017). Harnessing biodiesel-producing microbes: from genetic engineering of lipase to metabolic engineering of fatty acid biosynthetic pathway. Critical Reviews in Biotechnology, 37(1), 26–36.

26. Emmanuel, M. B., Evans, E. C., Abubakar, A., Labaran, L. M., Ali, A. V., & Zabe, M. (2020). Production, partial purification and characterization of lipase enzyme expressed by *Klebsiella pnemoniae* of vegetable oil contaminated soil. International Journal of Biochemistry and Biophysics, 8(2), 30–39.

27. Iqbal, A. S., & Rehman, A. (2015). Characterization of lipase from *Bacillus subtilis* I-4 and its potential use in oil contaminated wastewater. Brazilian Archives of Biology and Technology, 58, 789–797.

28. Mohan, T. S., Palavesam, A., & Immanvel, G. (2008). Isolation and characterization of lipase producing *Bacillus* strains from oil mill waste. African Journal of Biotechnology, 7, 2728–2735.

29. Jaiganesh, R., & Jaganathan, M. K. (2018). Isolation, purification, and characterization of lipase from *Bacillus sp*. From kitchen grease. Asian Journal of Pharmaceutical and Clinical Research, 11(6), 224–227.

30. Fathi, F., Kasra-Kermanshahi, R., Moosavi-Nejad, Z., & Qamsari, E. M. (2021). Partial purification, characterization and immobilization of a novel lipase from a native isolate of *Lactobacillus fermentum*. Iranian Journal of Microbiology, 13(6), 817–877.

31. Tripathi, R., Singh, J., Bharti, R. K., & Thakur, I. S. (2014). Isolation, purification and characterization of lipase from *Microbacterium sp*. and its application in biodiesel production. Energy Procedia, 54, 518–529.

32. Veerapagu, M., Narayanan, A. S., Ponmurugan, K., & Jeya, K. R. (2013). Screening selection identification production and optimization of bacterial lipase from oil spilled soil. Asian Journal of Pharmaceutical and Clinical Research, 6(3), 62–67.

